# Smart Parallel Automated Cryo Electron tomography

**DOI:** 10.1101/2023.12.14.571776

**Authors:** Fabian Eisenstein, Yoshiyuki Fukuda, Radostin Danev

## Abstract

*In situ* cryo electron tomography enables investigation of macromolecules in their native cellular environment. Data collection, however, requires an experienced operator and valuable microscope time to carefully select targets for tilt series acquisition. Here, we developed a workflow using machine learning approaches to fully automate the entire process, including lamella detection, biological feature segmentation, target selection, and tilt series acquisition, all without the need for human intervention.

## Main text

For decades, structural biologists have aimed to determine the structure of macromolecular complexes within their native cellular environments. Cryo electron tomography (cryoET) in conjunction with sample thinning methods, such as cryo-focused ion beam (cryoFIB) milling, has proven to be a powerful tool in this pursuit. However, obtaining structures at molecular resolutions similar to what is possible for purified complexes using single particle analysis (SPA) cryo electron microscopy (cryoEM) demands large quantities of data^1–5^. This is exacerbated by the scarcity and/or conformational heterogeneity of most macromolecules of interest within the cell.

Recently, much progress was made to streamline the cryoET workflow and improve throughput. Automated cryoFIB-milled lamella preparation^6–10^ and serial cryo lift-out^11,12^ provide an increasing number of samples for cryoET imaging. Beam-image shift cryoET enabled the collection of tens of tilt series on cryoFIB milled lamellae in parallel (PACEtomo), increasing the throughput from tens to hundreds of tomograms per day^16,17^. However, some bottlenecks remain.

*In-situ* cryoET samples have thicknesses of several hundred nanometres and therefore require high-end electron microscopes with high acceleration voltages and energy filters. While such instruments are becoming more common, usage fees remain high, and access time is limited for many researchers. Increasing the data collection throughput and minimising user input is thus highly desirable. In this regard, SPA cryoEM is far ahead of cryoET. Sample screening, area selection, and data acquisition have been mostly automated using a variety of deep learning approaches^13–15^, and while not widely adopted yet, a fully automated SPA cryoEM workflow is forthcoming. CryoET samples, however, are much more diverse than SPA samples and targeting areas of interest still requires operator experience. This makes the identification of hundreds of targets challenging and time consuming. It is common for operators to spend many hours on the microscope selecting targets instead of collecting data.

Here, we present the <u>s</u>mart <u>p</u>arallel <u>a</u>utomated <u>c</u>ryo <u>e</u>lectron <u>tomo</u>graphy (SPACEtomo) package, which uses machine learning to eliminate all manual steps in the cryoET data acquisition workflow and enables unattended acquisition of any sample loaded into the microscope.

The conventional setup for a cryoET microscope session, using a microscope control and automation software like SerialEM^18,19^, typically involves several steps (Fig. 1a). The sample grid is loaded into the microscope, low magnification montage (LM) maps are taken to identify lamellae and medium magnification montage (MM) maps are collected for each lamella individually. The MM maps are then inspected by the expert user, regions of interest are manually selected, and high magnification preview images are collected for the automated realignment procedure during batch tilt series collection. Finally, automatic data collection is started using either SerialEM’s tilt series controller for single tilt series or PACEtomo for parallel acquisition. The critical stage requiring the most operator time and expertise is the feature detection and target selection.

**Figure 1.**
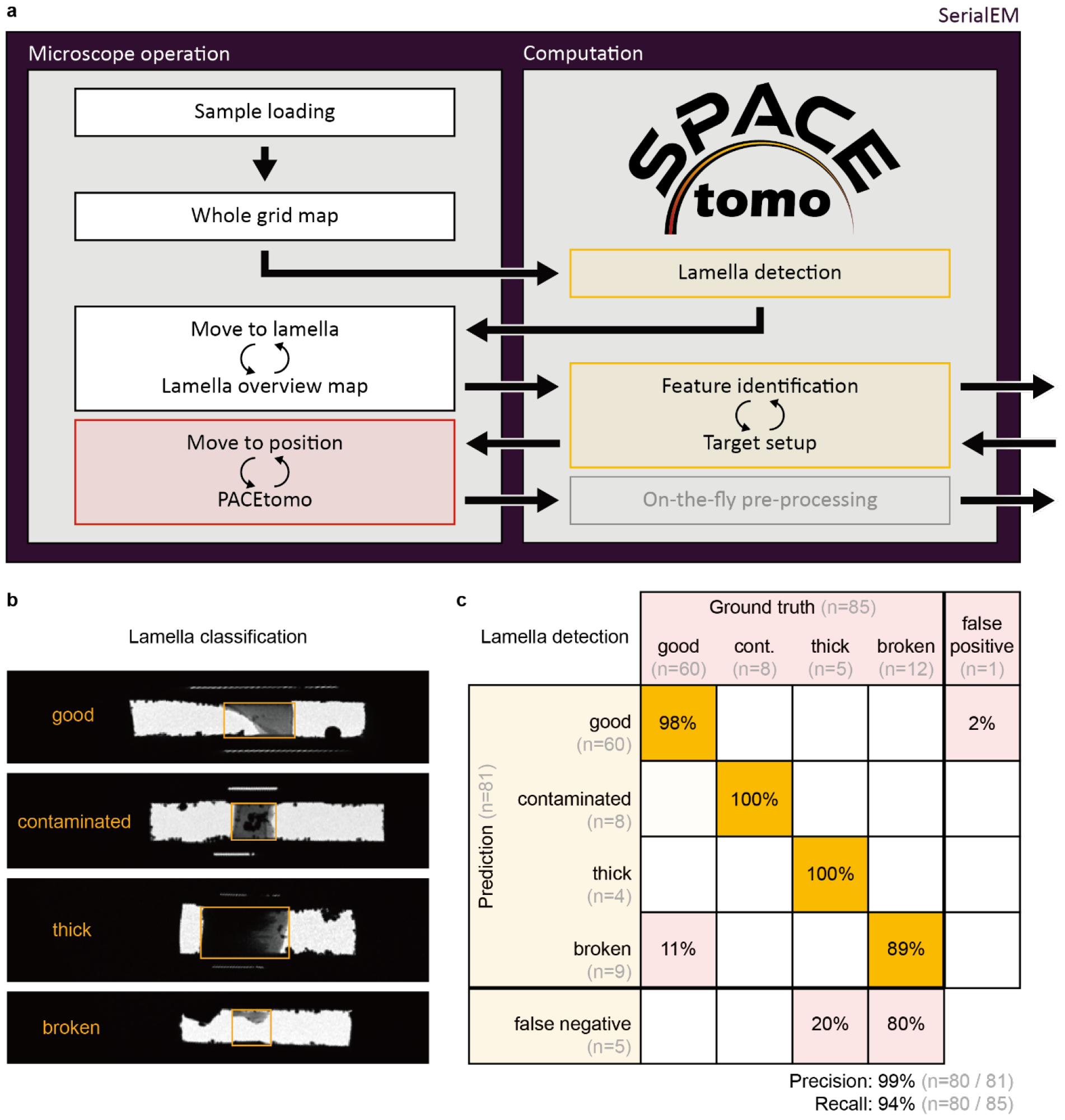
CryoET data acquisition workflow on *in situ* lamellae and automated lamella detection. a. Schematic showing the cryoET workflow. Yellow boxes can be automated using SPACEtomo. Red box is already automated by PACEtomo inside SerialEM. Arrows leaving the frame indicate options for external processing besides the microscope control software. b. Examples for each class of lamella in the lamella detection training dataset. c. Performance of the trained lamella detection model on a test dataset (n=85).

SPACEtomo addresses both, lamella localisation and target selection, by employing deep learning algorithms to analyse LM maps and MM maps of lamellae. These models can be deployed from within SerialEM or, in need of more GPU performance, can be run on an external processing workstation.

Lamella detection benefits from the characteristic shape and strong contrast of lamellae on commonly electron opaque samples, making it relatively robust. An out-of-the-box implementation of YOLOv8^20^ was trained on manually labelled LM maps to detect and classify lamellae into four classes (Fig. 1b). On a test dataset (n=85) the model achieved a detection rate (recall) of 94% (Fig. 1c). The bounding boxes determined by the model guide the acquisition position and dimensions of MM maps.

The collected MM maps serve as the basis for the identification of cellular structures, which is a challenging task due to low image contrast and substantial morphological variability. We utilised a modified U-Net^21^ for 2D segmentation of MM maps (Fig. 2a), leveraging its success in biomedical image segmentation. The model architecture and hyperparameters were configured in an automated manner by nnU-Net^22^. Training data was generated by manually segmenting MM maps of starved Yeast cells^23^ (*S. Cerevisiae*) lamellae and iteratively correcting predictions using a human-in-the-loop approach. The trained nnU-Net produced segmentations suitable for target selection for subsequent high-magnification tilt series collection (Fig. 2b).

**Figure 2.**
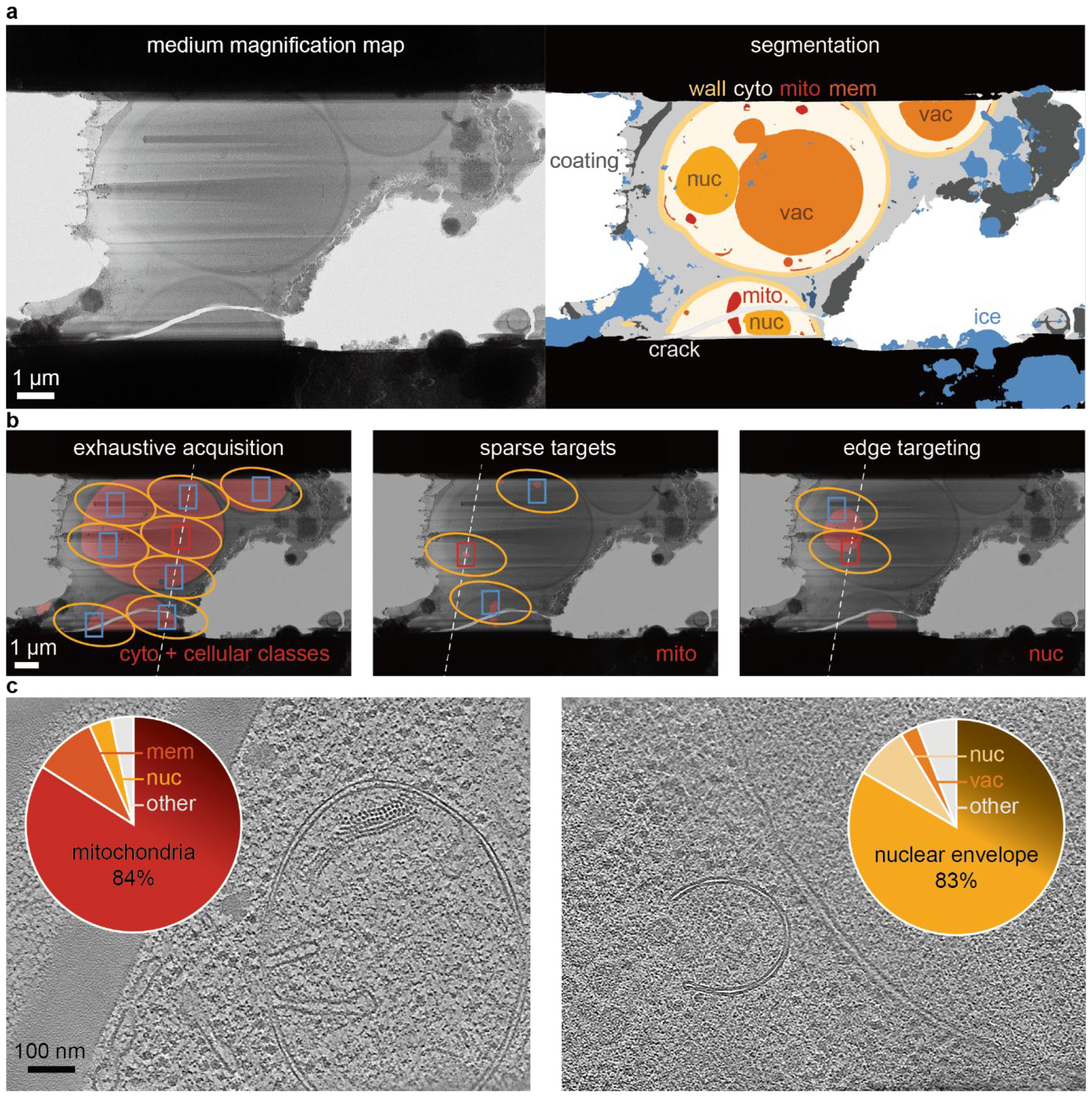
Feature detection and target selection. a. Representative MM map of a Yeast lamella (left) and its 2D segmentation (right). Yellows, oranges and reds show biological classes. Blues and greys show contamination and ice. A subset of the classes is labelled by abbreviations. Cyto: cytoplasm, wall: cell wall, nuc: nucleus, vac: vacuole, mito: mitochondria, mem: membranes, coating: lamella coating, ice: ice contamination, crack: crack in lamella. b. Examples for the three modes of target selection: Exhaustive acquisition, e.g. cellular area (left), sparse targets, e.g. mitochondria (middle) and edge targeting, e.g. nuclear envelope (right). Masks of the corresponding classes of interest are shown as a red overlay. Beam shape at maximum tilt angle is marked by yellow ellipse. Camera fields of view are outlined in red for the tracking target and blue for other targets. The dashed white line represents the tilt axis orientation. c. Slices through examples of acquired tomograms using SPACEtomo targeting mitochondria (left) and the nuclear envelope (right). Pie charts show the fraction of tilt series containing the target (n=299 and n=187, respectively) as well as common false positives (mem: cytoplasmic membranes, nuc: nucleus/nucleoplasm, vac: vacuole).

Target selection for *in situ* cryoET is highly dependent on the specific biological focus. To accommodate diverse use cases, we implemented three distinct approaches to target selection guided by a target mask based on the lamella segmentation (Fig. 2b). Option 1 is maximum coverage or exhaustive acquisition. A hexagonal grid of target positions spaced to minimize exposure overlap according to beam diameter and maximum tilt angle is generated, positions falling outside the target mask are removed and the remaining points are optimized for coverage. Option 2 places targets arbitrarily within the target mask and positions are optimised for coverage in clusters. This approach is useful for sparsely distributed targets like mitochondria. Finally, option 3 allows aiming for area boundaries rather than optimising coverage, for example when aiming for the nuclear envelope. While these target selection options should cover most use cases, adjustment and implementation of new targeting schemes are straightforward since the procedure is independent of the automation and deep learning steps.

To facilitate the adoption of SPACEtomo for new users, we implemented all steps in a set of Python scripts that can be executed within SerialEM, necessitating no additional software or hardware beyond that of conventional cryoET approaches. After loading sample grids into the microscope and ensuring microscope alignment, a specific biological target, such as mitochondria or the nucleus, can be set and the SPACEtomo script started. It will carry out the session autonomously and the result will be the collected tilt series. Utilising external workflows and software for automated on-the-fly processing^24–26^, the entire cryoET workflow can be operated seamlessly from sample loading to tomogram generation, requiring no human intervention.

We used SPACEtomo to collect 299 tilt series on 29 lamellae of cryoFIB milled Yeast cells targeting mitochondria (Fig. 2c, left, Table S1). 84 % of the resulting tilt series contained whole or partial mitochondria. An additional experiment targeting the nuclear envelope on 34 lamellae resulted in 187 tilt series of which 83 % included segments of nuclear envelope and an additional 8 % showed nucleoplasm (Fig. 2c, right, Table S2).

These results prove that the future of cryoET data collection lies in complete automation. SPACEtomo provides the first workflow without any human intervention making use of deep learning models to detect targets on the sample grid and setup parallel data acquisition. The result is a substantial reduction in both operator time and microscope idle time leading to greater productivity.

While the current segmentation model is trained on *S. cerevisiae* lamellae, extending it to other organisms is a straightforward process when generating training data using a human-in-the-loop approach. General classes such as “ice”, “crack” and “coating” are universally applicable and already transferable to other samples when aiming for exhaustive acquisition of lamellae (Fig. S4 and S5).

Some operators will be hesitant to entrust target selection to a machine learning model. In such cases, the automated workflow can be run up to a desired level, allowing for manual inspection and selection/modification of targets using the PACEtomo scripts.

As models continue to improve and more and more organisms will be covered, fully automated cryoET will become even more robust and commonplace in the field of *in situ* structural biology.

## Acknowledgements

We are grateful to Anna Bieber and Cristina Capitanio for providing the Yeast lamella montages used to create the training dataset. Additionally, we thank Matthias Pöge, Gregor Weiss and Sven Klumpe for providing training data for the lamella detection model, and Hiroka Kashihara and Sachiko Tsukita for providing the EpH4 cell sample. We express our appreciation for fruitful discussions about deep learning approaches with Sven Klumpe, Yusuke Hirabayashi, Shogo Suga and Hiroki Kawai. We extend our gratitude to Masahide Kikkawa for providing feedback and support. Further thanks go to Yoichi Sakamaki, Tony Wang, and Kazuhiko Nakamura for the support of the Graduate School of Medicine cryoEM facility. F.E. was an International Research Fellow of the Japan Society for the Promotion of Science (JSPS, #P20764) and was recipient of a Grant-in-Aid for Scientific Research (KAKENHI, 21F20764). Y.F. (KAKENHI, JP19H05707) and R.D. (KAKENHI, 22H02554) were supported by JSPS grants.

## Author contributions

F.E. developed SPACEtomo, labelled the training data and conducted all cryoFIB and cryoET experiments. Y.F. prepared the Yeast samples and helped with troubleshooting. F.E. and R.D. designed the experiments. F.E. wrote the manuscript with contributions from all authors.

Correspondence to

## Methods

### Lamella detection

An out-of-the-box implementation of YOLOv8^20^ was trained on 614 LM map pieces of varying pixel sizes (rescaled to 400 nm/pixel) containing 1076 examples of lamellae. Lamellae were classified into “good”, “contaminated”, “thick” and “broken” lamellae (Fig. 2a). To enhance rotational invariance, the 614 LM map pieces were added to the training dataset again with flipped axes. An additional 534 LM map pieces not containing any lamellae were added to the training dataset.

For the practical analysis of a whole montage, the LM map is split into subregions and analysed by the model (Fig. 1b). Resulting bounding boxes are checked for overlap and merged accordingly to obtain a single bounding box per lamella. Lamellae are also classified into 4 categories. A user may choose to exclude any particular class for subsequent analysis, but in this work, only lamellae categorised as “broken” were excluded. Using the coordinates determined by the lamella identification model, SerialEM navigator points are created, and MM maps are acquired at all lamella positions. Due to magnification offsets in the coordinate system, an additional image at an intermediate magnification can be taken to refine the position of the detected lamella and ensure that the MM map is centred accordingly. To obtain MM maps with sufficient contrast, defocus values between 50 and 100 micrometres were used at a pixel size of 1.89 nm per pixel.

### Lamella segmentation

The model architecture and hyperparameters were configured in an automated manner by nnU-Net^22^. Only the patch size was adjusted by a factor of 2 manually to allow the model to consider more context. Generation of training data is generally a bottleneck for machine learning approaches. Here, we used a human-in-the-loop approach to speed up labelling. Initially, 10 MM map pieces from starved *S. cerevisiae*^23^ were manually segmented into 17 categories (including several background categories, Fig. S2) and used to train a 2D nnU-Net. The trained nnU-Net provided imperfect segmentations of 10 additional MM map pieces that were manually corrected and added to the training dataset. This was repeated until the training dataset contained 50 images. Ten images of Baker’s Yeast under exponential growth conditions (as described below) were added to enhance performance. Five-fold cross-validation as implemented by nnU-Net was applied during training and ensemble inference was used for the lamella segmentation to further improve predictions. Segmentations were not post-processed. MM maps were minimally pre-processed by scaling to the model pixel size of 2.28 nm per pixel and exporting as PNG file.

MM map segmentation is the most computationally expensive step of the workflow. It requires from several minutes to several tens of minutes on a typical Gatan K3 control computer equipped with a Nvidia Quadro P6000 (Fig. S3). We outsourced the inference step to a separate script that is automatically called from the main SPACEtomo script in SerialEM upon collection of a new MM map. Thus, the analysis can run in the background, while the microscope can collect the next montage or set up targets on previously analysed lamellae. Additionally, the lamella detection model can be utilised to determine the bounding box of the lamella within the MM map. This can be achieved by downscaling the map to the pixel size of the lamella detection model and padding it to the minimum input size before running the inference. The resulting bounding box can be used to exclude anything outside the lamella area from the segmentation procedure. Finally, we implemented an option to save the MM maps in a separate folder that can be monitored by the *SPACEtomo_monitor*.*py* script, which must be run separately on a GPU machine or cluster that has access to the folder. This script will detect any new MM map files and queue them up for analysis. While running an external machine should generally not be necessary, it can be done to eliminate any computational bottlenecks (Fig. S3).

### Target setup

Independent of the modality of target selection, the area of interest is generated by combining all segmented classes in a user given target list to a single target mask. Furthermore, the user can specify a list of classes to avoid. The segmentation of these classes will be combined to a penalty mask affecting the score function and hence, the target placement.

For maximum coverage, a hexagonal grid of target positions spaced according to the beam diameter and maximum tilt angle is generated. Target positions that fall out of the area of interest are removed and the remaining points are translated for maximum coverage using a stochastic gradient descent (SGD) algorithm to maximise a score function based on how much of the field of view is covered by the area of interest. This is done iteratively removing targets that fall below the score threshold at every step.

For sparse target selection, points are placed wherever the segmentation shows an area of interest. The points are then grouped into clusters based on distance to each other to avoid overlapping exposures. These clusters are optimised for coverage independently using SGD and a weighted score function to encourage placement of a small target area in the centre of a tilt series. Between iterations, targets are excluded based on score threshold and distance to neighbouring targets. The point clustering is reevaluated at every iteration. Upon convergence, an additional step checks if any adjacent points can be placed to cover additional area of interest, which would initiate an additional iteration of refinement.

Targeting edges of areas of interest works analogously to the sparse target setup. Instead of optimising coverage, the score function was adjusted to optimise for covering half of the acquisition area by the target.

To choose the tracking area for PACEtomo, one of the targets is selected automatically depending on the distance from the centre and a score based on the penalty mask alone. Since the tracking target could be compromised by dark areas or ice coming into the field of view at higher tilt angles, this score considers the camera field of view at the maximum tilt angle.

SPACEtomo will further attempt to find points to measure the sample geometry using SerialEM’s beam tilt autofocus as implemented in PACEtomo. For this purpose, it will use the maximum coverage target setup to search for positions that are neither in the target nor in the penalty mask. If less than three such points can be found it will default to using the sample geometry values set by the user in the PACEtomo script.

### Yeast sample preparation

Baker’s yeast obtained from a local supermarket was grown in filtered YPD medium (1% yeast extract, 2% peptone, and 2% glucose) at 30°C and shaking at 200 rpm overnight. Before plunge freezing, 0.5 mL culture (OD_600_=3.49) was spun down at 2000x g and resuspended in 250 or 125 µL YPD containing cryoprotectant (15 % dextran and 2 % glucose).

The sample was applied to glow discharged Quantifoil Cu200 R1/4 grids, mounted in a Vitrobot Mark IV (Thermo Scientific) and back blotted for 12-15 s between a Teflon sheet and a filter paper, before plunging in ethane/propane mixture.

### CryoFIB milling

Sample grids were clipped in modified autogrids (Thermo Fisher Scientific) allowing for lower incidence angles of the focused ion beam. An Aquilos 2 dual-beam instrument (Thermo Fisher Scientific) was used for the cryoFIB milling process. After sputter-coating with Pt for 15 s at 30 mA and organometallic Pt coating for 20 s using the integrated gas injection system, clumps of yeast cells were identified manually using MAPS 3.21 (Thermo Fisher Scientific). Automated preparation, milling and polishing down to a target thickness of 200 nm was done in AutoTEM Cryo 2.4 (Thermo Fisher Scientific) reducing the ion beam current stepwise from 1 nA to 30 pA. Between 10 and 25 lamellae were targeted per grid. Samples were stored under liquid nitrogen until data collection.

### Data collection

Tilt series were collected on a Thermo Scientific Krios G4 transmission electron microscope (Thermo Fisher Scientific) operated at 300 kV and equipped with a BioQuantum energy filter and K3 direct electron detector (Gatan). The microscope and camera were controlled by SerialEM 4.1.0beta^18,19^. SPACEtomo was used for automated mapping and target setup with the setting shown in Table S3. The “target_score_threshold”, in particular, can be used to optimize for precision or recall during target selection (Fig. S6). SPACEtomo was run four times on three grids, twice targeting mitochondria, and twice targeting the nuclear envelope. Hence, one of the grids was imaged twice using different targets. PACEtomo was automatically called for dose-symmetric tilt series acquisition covering 120° with 3° increments centred on a start tilt angle of -9° to partially compensate for lamella pretilts. A total electron fluence of 178.4 e^-^/Å^2^ and a defocus of 5 µm was targeted (Table S4).

### Tomogram reconstruction

Tilt series were aligned and reconstructed using AreTomo2^25^ and denoised using cryo-CARE^27^. IMOD^28,29^ was used for tomogram visualisation. Tomograms and tilt series were manually inspected to confirm presence of target structures.

### Implementation details

SPACEtomo was implemented in Python 3.9 using the deep leaning frameworks nnUNetv2 and Ultralytics. Additional Python libraries include: PyTorch, mrcfile, Matplotlib, NumPy, Pillow, scikit-image, scikit-learn and SciPy.

## Data availability

Lamella montages and raw tilt series frames produced during SPACEtomo development were deposited in the Electron Microscopy Public Image Archive under accession codes EMPIAR-XXXXX.

Labelled training datasets for deep learning models are available at https://doi.org/10.5281/zenodo.10360315 and https://doi.org/10.5281/zenodo.10360344.

Trained deep learning models were deposited at https://doi.org/10.5281/zenodo.10360489 and https://doi.org/10.5281/zenodo.10360540.

## Code availability

All SPACEtomo Python scripts are available at https://github.com/eisfabian/SPACEtomo. SPACEtomo is currently still in an alpha stage and will be updated to improve performance and reliability and add additional features and models.

## Supplementary material

**Figure S1.**
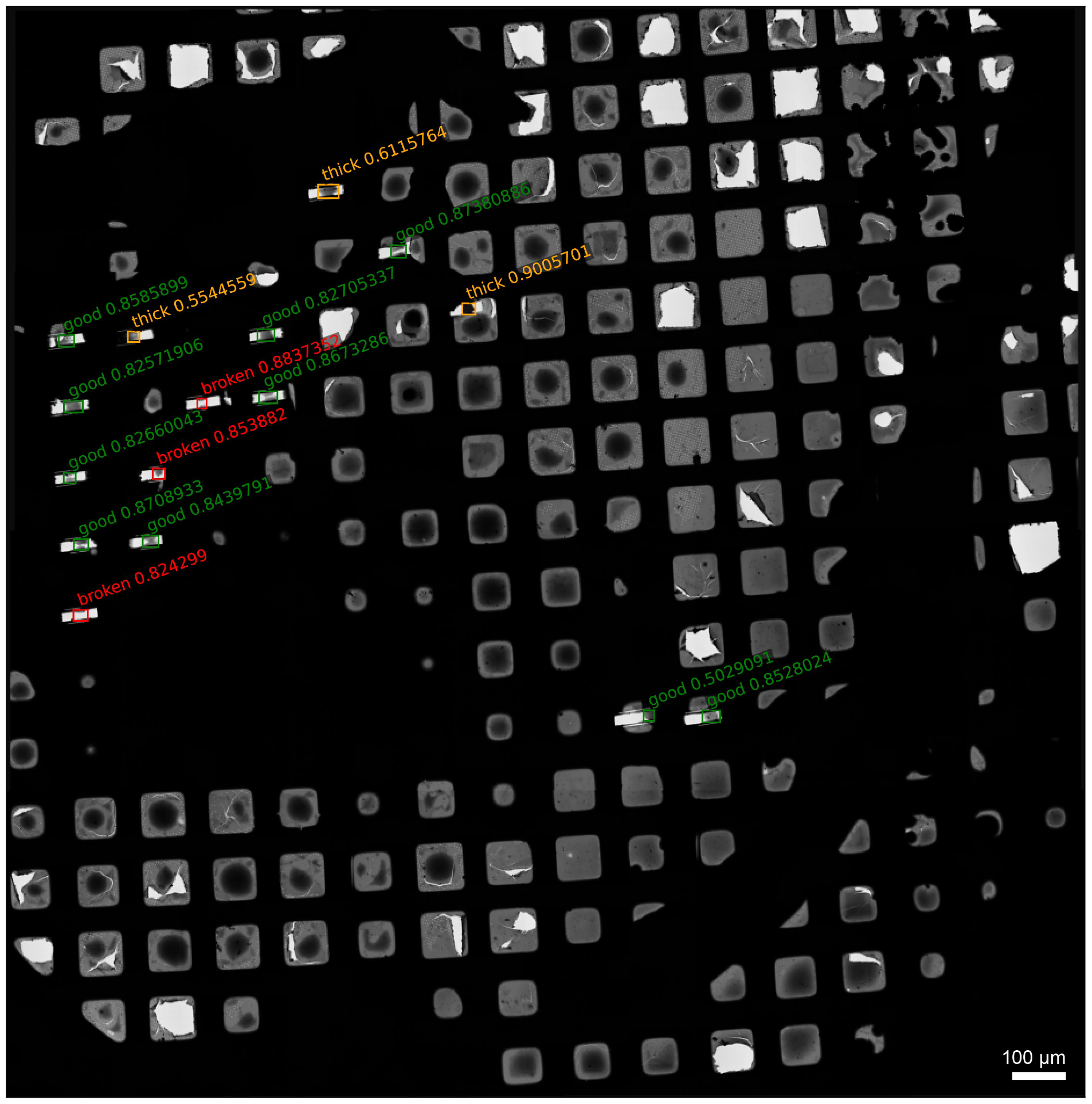
Example of lamella detection on a whole grid montage LM map. Shown is a cryoFIB-milled grid of Yeast cells with 16 detected lamellae. Bounding boxes of detected lamellae are coloured according to their predicted class (green: “good”, red: “broken”, yellow: “thick”). The label for each bounding box shows the class label and the confidence score of the model.

**Figure S2.**
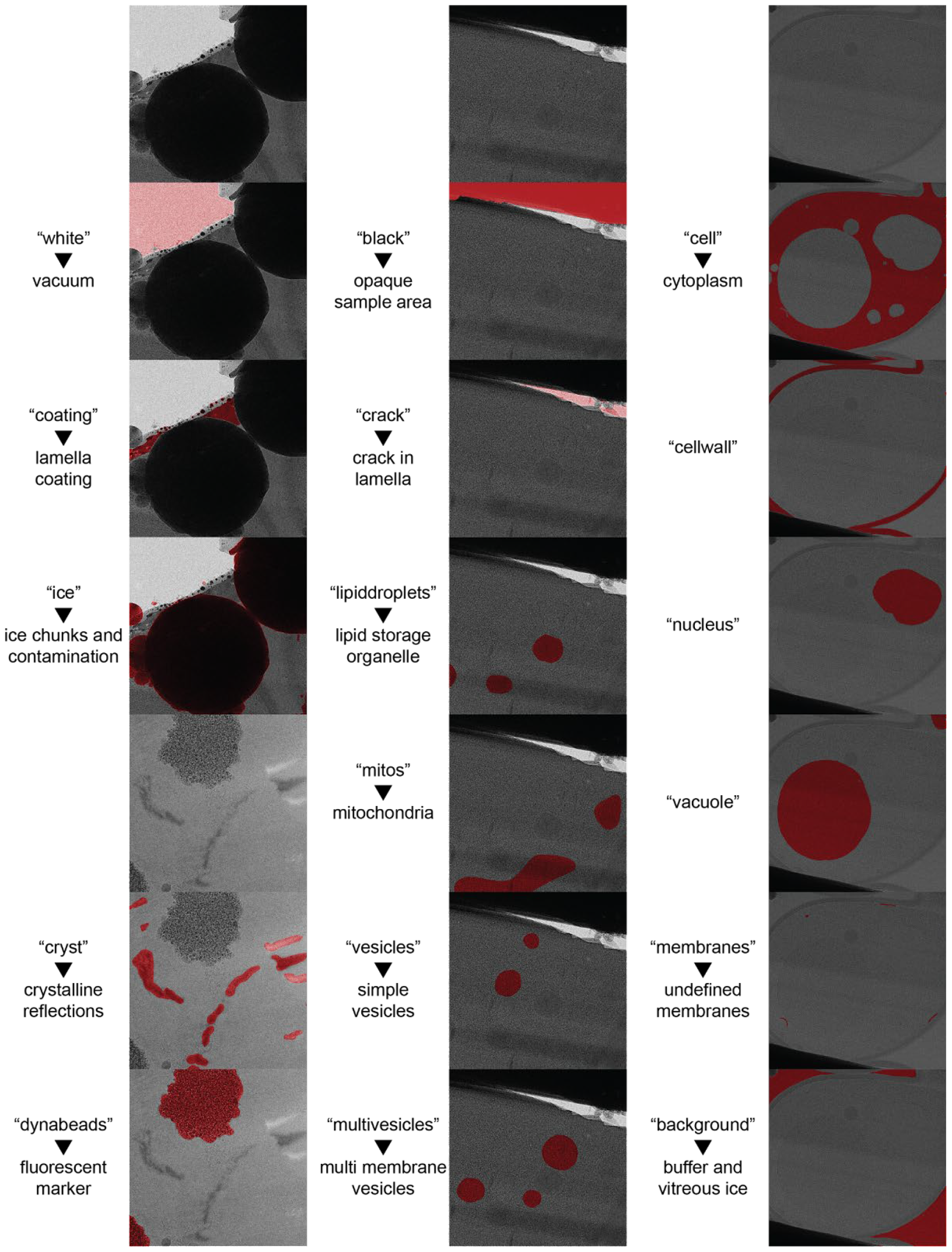
Segmentation classes. Examples for all 17 classes used to train the nnU-Net for lamella segmentation. Class names in quotes are accompanied by short descriptions, where necessary, and an example image showing the respective class segmentation in red.

**Figure S3.**
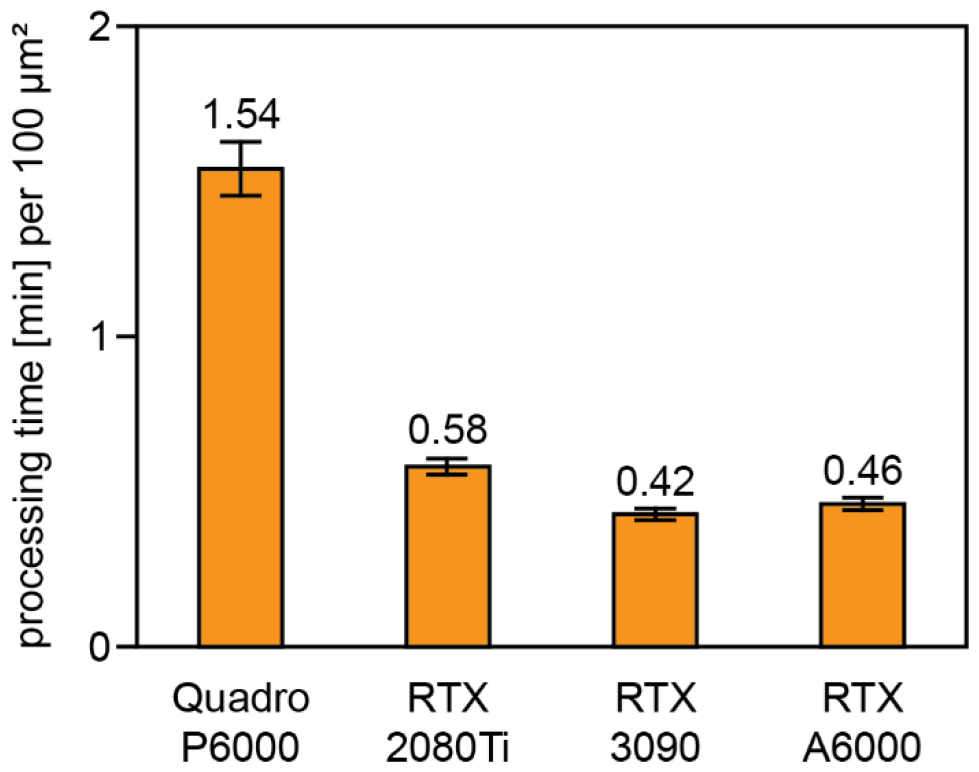
Segmentation inference time on different GPUs. Mean segmentation inference time in minutes per 100 µm^2^ obtained from inferences of 8 MM maps of sizes between 300 and 1000 µm^2^. Error bars show the standard deviation.

**Figure S4.**
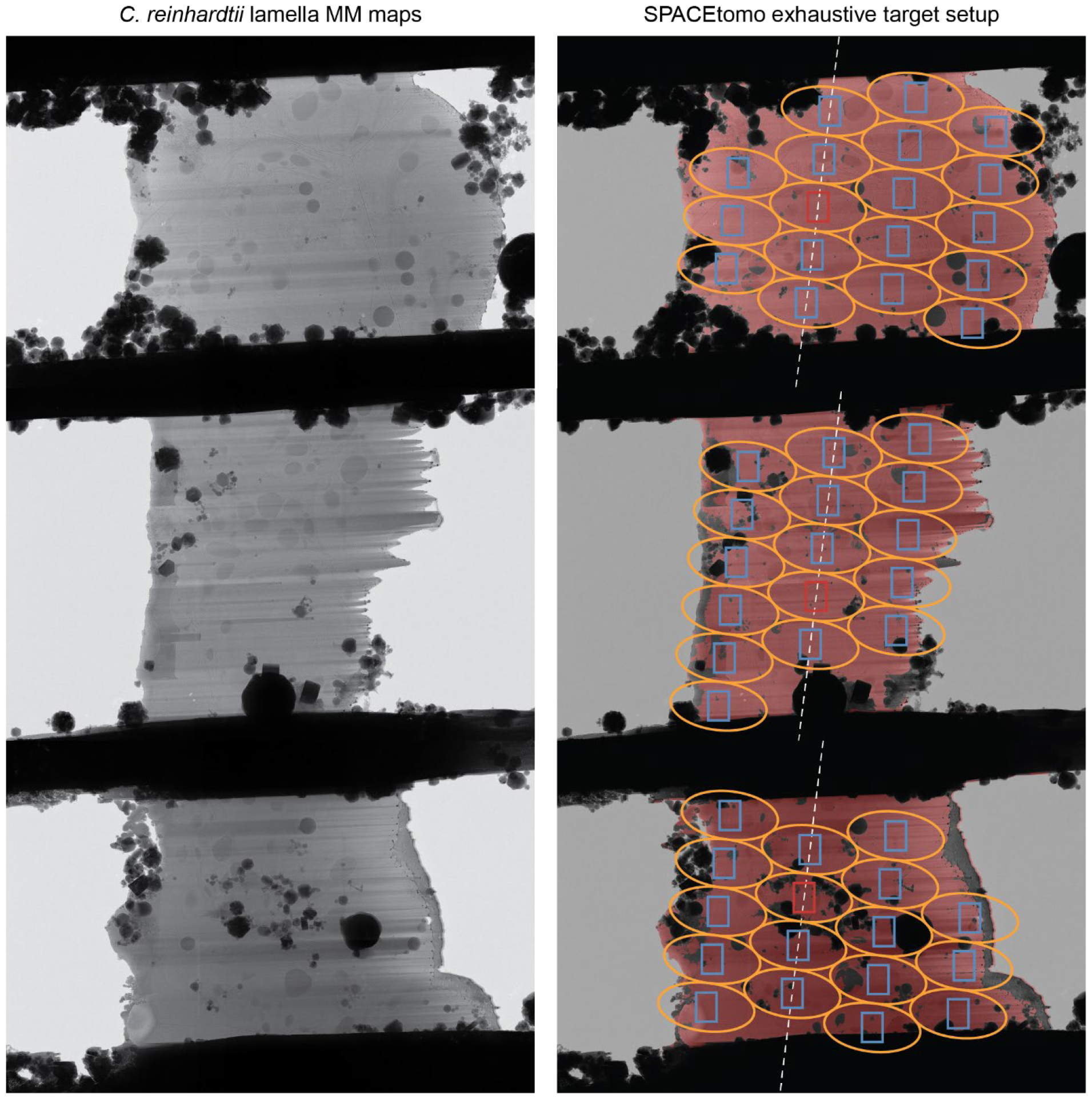
Transferability of the SPACEtomo segmentation model to *C. reinhardtii*. Shown are 3 examples of *C. reinhardtii* lamella MM maps on the left and the automated SPACEtomo target selection giving a target list of all classes except the organism independent classes to be avoided (e.g., black, white, ice, crack). Beam shape at maximum tilt angle is marked by yellow ellipse. Camera fields of view are outlined in red for the tracking target and blue for other targets. The dashed white line represents the tilt axis orientation.

**Figure S5.**
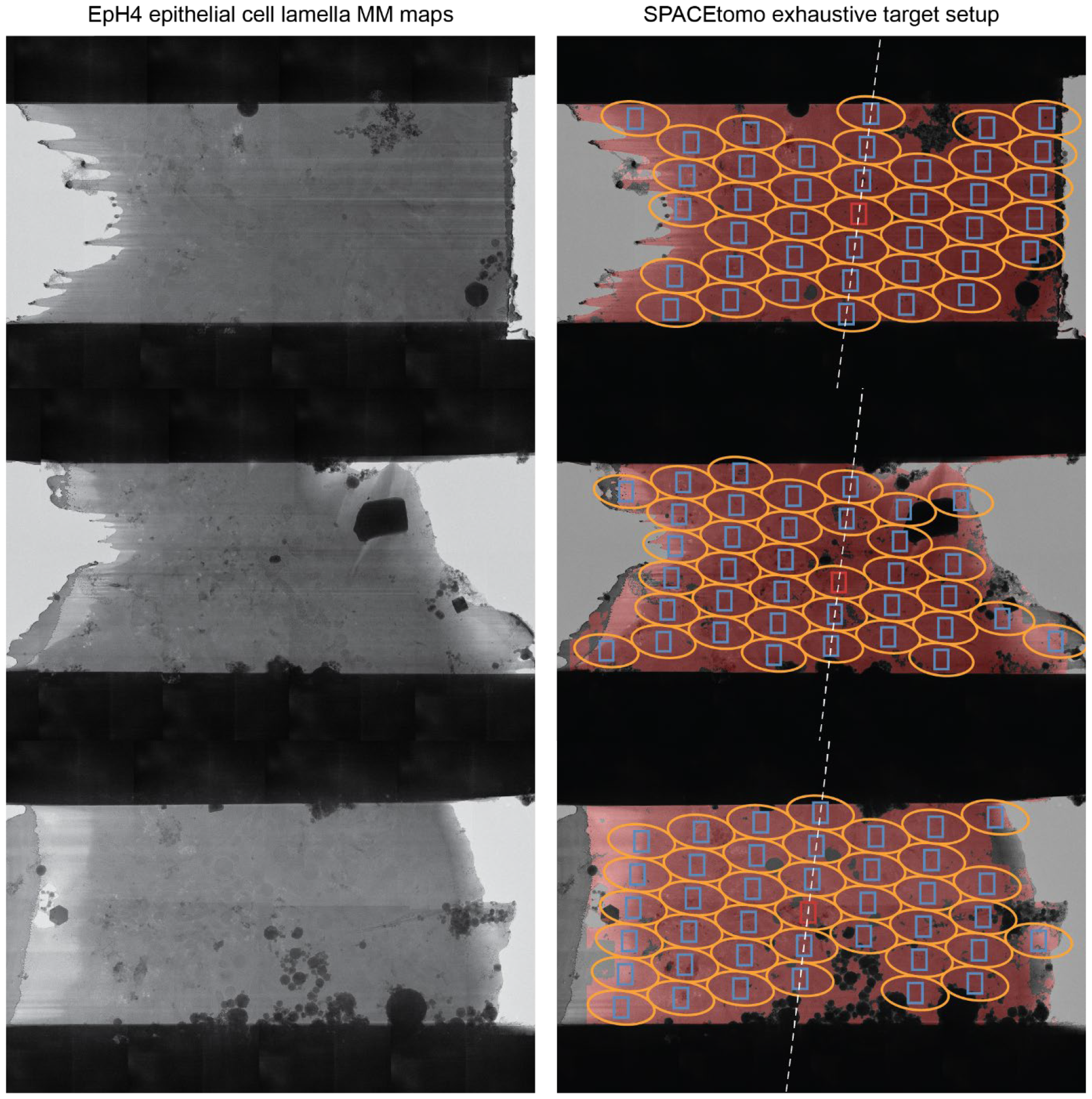
Transferability of the SPACEtomo segmentation model to EpH4 mouse epithelial cells. Shown are 3 examples of EpH4 cell lamella MM maps on the left and the automated SPACEtomo target selection giving a target list of all classes except the organism independent classes to be avoided (e.g., black, white, ice, crack). Beam shape at maximum tilt angle is marked by yellow ellipse. Camera fields of view are outlined in red for the tracking target and blue for other targets. The dashed white line represents the tilt axis orientation.

**Figure S6.**
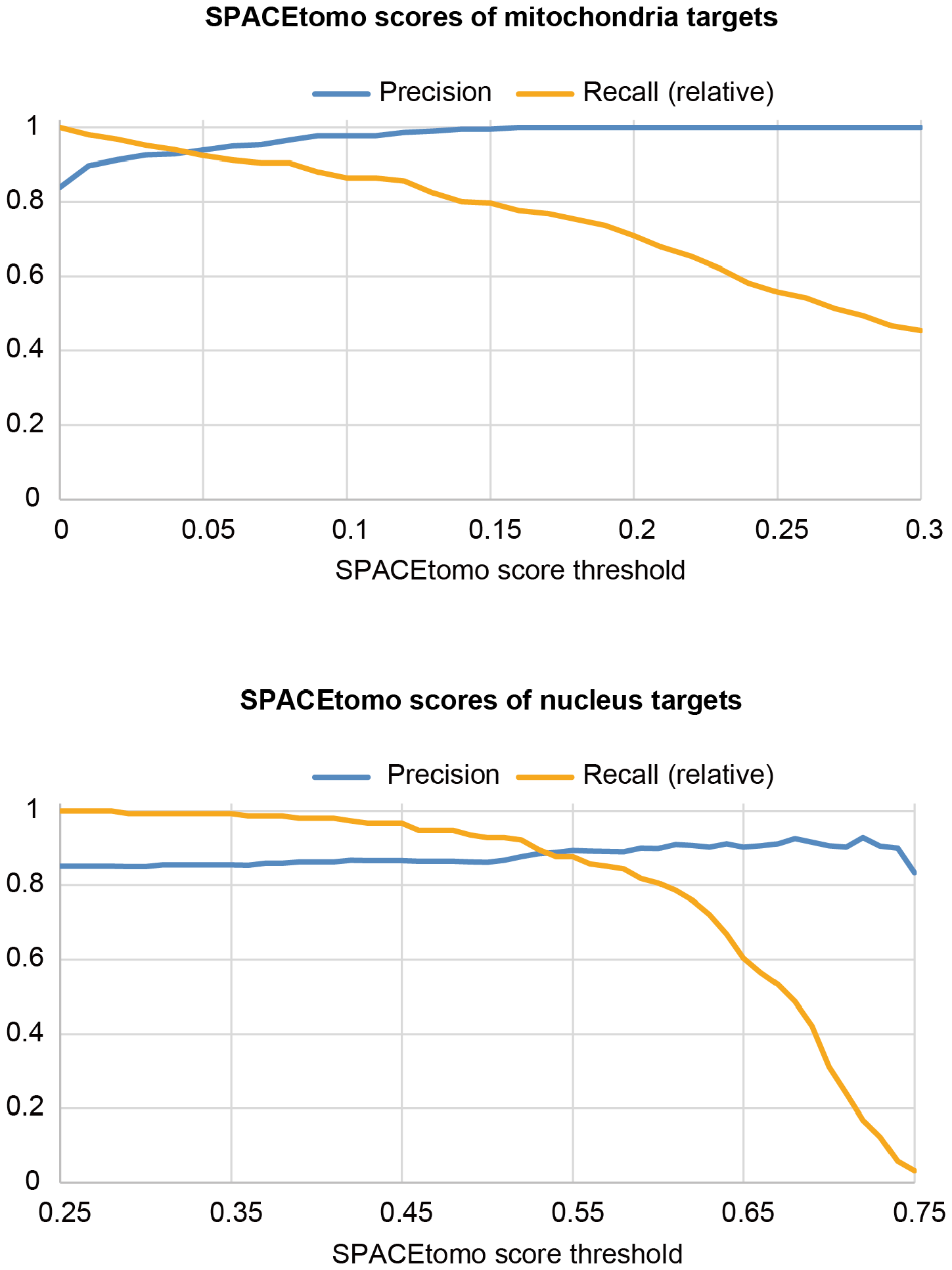
The SPACEtomo score threshold can improve precision of targets. Top: Precision and recall relative to used threshold (0.0) for 299 tilt series collected targeting mitochondria. Setting the threshold to 0.01 would have raised the precision to 90 % while losing only 2 % of mitochondria targets. Bottom: Precision and recall relative to used threshold (0.25) for 187 tilt series collected targeting the nuclear envelope. Even high thresholds did not yield a precision of 100 % indicating the presence of misclassifications in the segmentations that could be improved with more training data.

**Table S1:** SPACEtomo data collection targeting mitochondria.

|  | Mitochondria | Membranes | Nucleus | Other | Total |
| --- | --- | --- | --- | --- | --- |
| Tomograms # | 251 | 28 | 10 | 10 | 299 |
| Fraction [%] | 83.9 | 9.4 | 3.3 | 3.3 | 100 |

**Table S2:** SPACEtomo data collection targeting the nuclear envelope.

|  | Nuclear envelope | Nucleoplasm | Vacuole | Other | Total |
| --- | --- | --- | --- | --- | --- |
| Tomograms # | 156 | 15 | 5 | 11 | 187 |
| Fraction [%] | 83.4 | 8.0 | 2.7 | 5.9 | 100 |

**Table S3:**
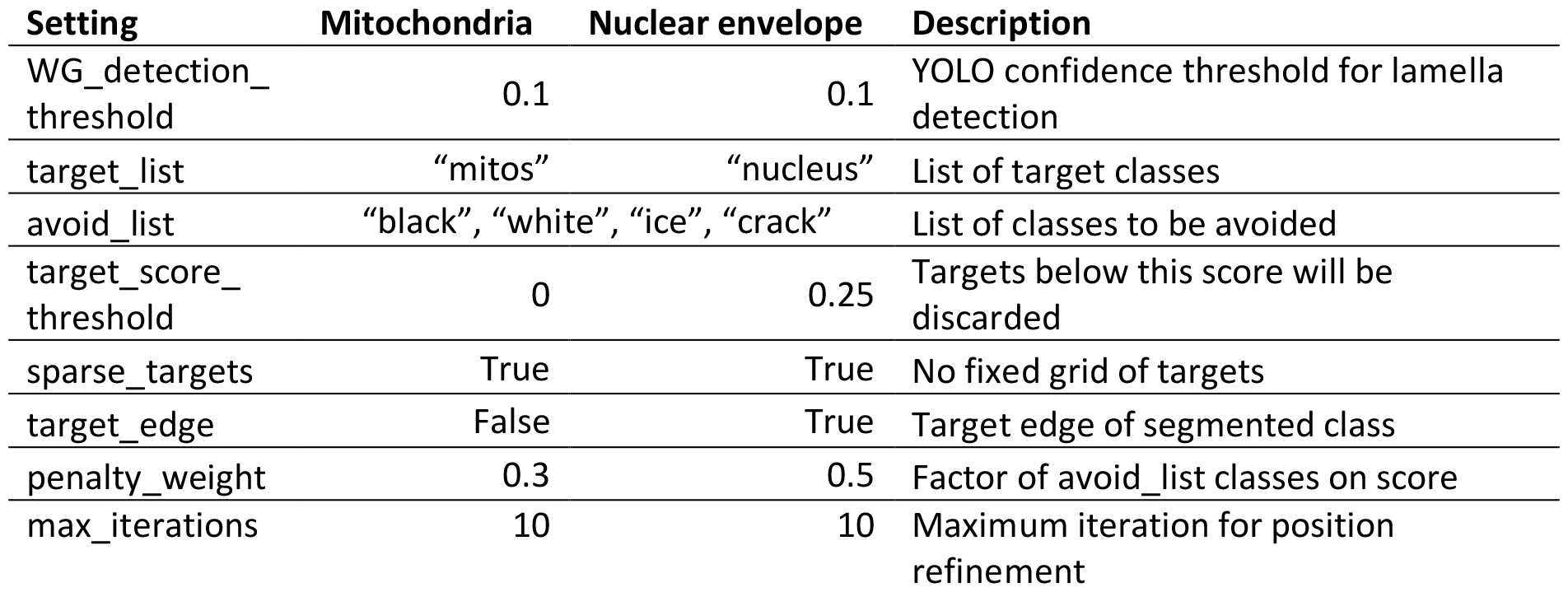
SPACEtomo target selection settings.

**Table S4:** Data collection parameters.

|  |  |
| --- | --- |
| Microscope | Krios G4 |
| Voltage [kV] | 300 |
| C2 aperture [μm] | 50 |
| Objective aperture [μm] | 70 |
| Electron detector | Gatan K3 |
| Energy filter slit width [eV] | 20 |
| Pixel size [Å] | 2.09 |
| Electron flux [e-/pixel/s] | 19 |
| Exposure time per tilt [s] | 1 |
| Movie frames per tilt | 10 |
| Electron fluence per tilt (e-/Å<sup>2</sup>) | 4.35 |
| Total electron fluence (e-/Å<sup>2</sup>) | 178.4 |
| Tilt range (°) | -51 to 69 |
| Tilt increment (°) | 3 |
| Target defocus (μm) | -5 |

